# Phage DisCo: targeted discovery of bacteriophages by co-culture

**DOI:** 10.1101/2024.11.22.624878

**Authors:** Eleanor A. Rand, Siân V. Owen, Natalia Quinones-Olvera, Kesther D. C. Jean, Carmen Hernandez-Perez, Michael Baym

## Abstract

Phages interact with many components of bacterial physiology from the surface to the cytoplasm. Although there are methods to determine the receptors and intracellular systems a specified phage interacts with retroactively, finding a phage that interacts with a chosen piece of bacterial physiology *a priori* is very challenging. Variation in phage plaque morphology does not to reliably distinguish distinct phages and therefore many potentially redundant phages may need to be isolated, purified, and individually characterized to find phages of interest. Here, we present a method in which multiple bacterial strains are co-cultured on the same screening plate to add an extra dimension to plaque morphology data. In this method, Phage *Dis*covery by *Co*-culture (Phage DisCo), strains are isogenic except for fluorescent tags and one perturbation expected to impact phage infection. Differential plaquing on the strains is easily detectable by fluorescent signal and implies that the perturbation made to the surviving strain in a plaque prevents phage infection. We validate the phage DisCo method by showing that characterized phages have the expected plaque morphology on Phage DisCo plates and demonstrate the power of Phage DisCo for multiple targeted discovery applications, from receptors to phage defense systems.

## Introduction

Bacteriophages (phages) are extremely abundant, diverse, and under sampled in the environment [1]. There are an estimated 10^31^ phages on Earth [2,3], and, beyond their plentiful number, phages may represent some of the largest reservoirs of unexplored genetic diversity on the planet [4]. Although this wealth of diversity can be sampled in almost any environment from soil to the ocean to the human gut [1], the distribution of phage diversity is not predicted to be uniform across environments [5]. Instead, as has been recognized by ecologists since the 1940s [6], sequence-based analysis suggests there are huge differences in the abundance of different groups of phages, with a relatively large percentage of the phage population made up of highly abundant, similar phages while most diversity is found in low abundance, rare phages [7]. Current culture-based methods for phage discovery are intrinsically biased towards the abundant majority and therefore increased phage discovery efforts in the absence of methodological innovation will inevitably lead to over-sampling of abundant phages, with only small and random glimpses at rare, phenotypically diverse phage populations.

Technologies have advanced to explore the characteristics of a given phage once it has been isolated in culture, for example receptor determination [8] or gene essentiality [9], while the opposite problem, finding a phage that uses a specific receptor or with other specific characteristics from an environmental sample, remains a laborious task. Characteristics of interest may include interaction with or selection against a bacterial receptor protein of interest [10–13], finding a phage that interacts with a putative defense system [14], selecting a diverse cocktail of phages [15], or other similar quests. Methods to isolate phages with desired characteristics have not advanced significantly beyond the original plaque assay methodology [16]. To find a phage of interest, a large panel of phages must be isolated on a single bacterial host strain, purified, replicated, and then screened retroactively on a range of bacterial hosts to identify candidates with various properties. The rarer the phage of interest, the more phages need to be isolated, processed, and screened.

Here, we describe a modification to the traditional plaque assay that allows for quick and efficient targeted isolation of environmental phages. We call this method Phage *Dis*covery by *Co-*culture (Phage DisCo). Phage DisCo works by culturing multiple bacterial strains in the same agar lawn and screening for differential plaquing via a fluorescence read-out. While we previously demonstrated that this approach allows for the direct isolation of plasmid-dependent phages [17], and fluorescence has been used to measure differential phage susceptibility between modified bacterial strains [18], here, we demonstrate its broader utility for general screening applications. With the ability to highlight differential lysis, we first isolate phages dependent on common *E. coli* phage receptors, followed by the targeted isolation of phages that interact with recently-described phage defense systems. Given the generality of the method, we anticipate Phage DisCo to be readily adaptable to a wide range of targeted phage discovery applications.

## Results

### Phage Discovery by Co-culture

Because phage diversity in the environment is unlikely to be uniformly distributed, naive phage isolation will repeatedly recover the most common phages while only occasionally recovering rare phages. This is illustrated by the classically modeled rank abundance curve (Figure 1A) showing that the most abundant phages make up a large proportion of the population while the bulk of the diversity is in the long tail of rare phages. Though phage plaque morphology offers a crude way to distinguish between different phages [19] it can be inconsistent, and is associated with a small number of variables making it an unreliable metric of diversity. In effect, current culture-based isolation methods are only able to sample the phage *population*, as opposed to sampling phage *diversity*.

**Figure 1:**
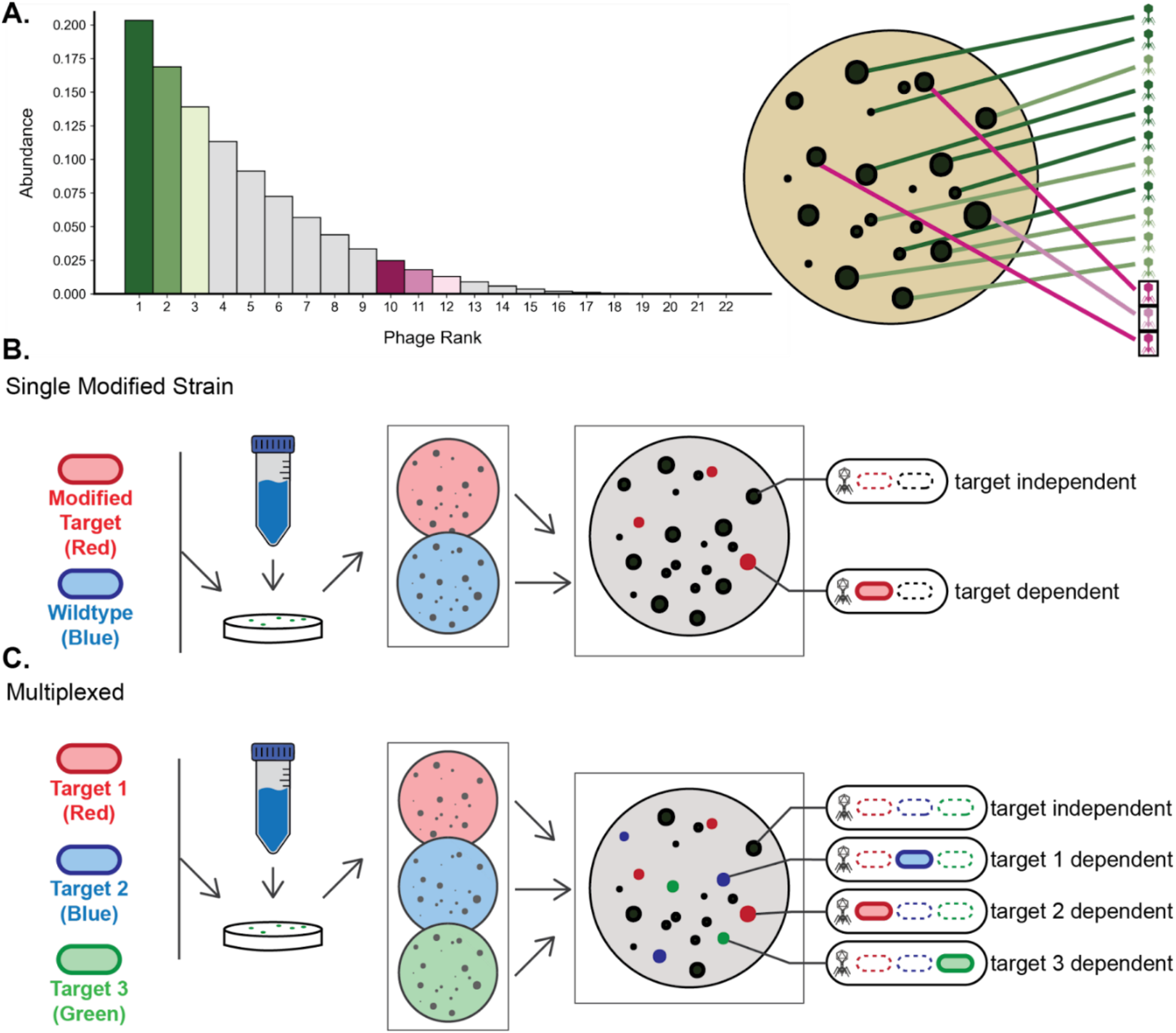
Phage Discovery by Co-culture (DisCo) allows for rapid identification of phages with properties of interest. **A**. Simulated rank/abundance curve illustrating that the diversity of phages in an environment at any given time is not expected to be evenly distributed across phage taxa. **B**. Conceptual diagram of the most simple co-culture screening set up. In this case, a wildtype strain and a singly modified strain are each tagged with a unique fluorescent protein and plated together with an environmental sample. After imaging in the individual channels and creating a composite image, differential plating can be detected. **C**. Conceptual diagram of a multiplexed co-culture system where three strains, each with a modification and a fluorescent tag are plated together with an environmental sample. This allows for extra dimensionality on each screening plate.

To improve the efficiency of targeted phage discovery, we devised a co-culture screening approach which effectively adds an additional dimension to traditional plaque assay data. In its simplest form, the Phage DisCo assay uses two strains: a wildtype and a modified strain, each with their own fluorescent tag. The modified strain can be altered in any way that might interact with phage infection, for example, by knocking out or modifying a phage receptor or by introducing a known or putative phage defense system. The wildtype and modified strain are then plated together with an environmental sample containing unknown phages. After growth, the plate is imaged in both fluorescent channels, and plaques with differential fluorescence between the two strains are identified, most readily by a composite image (Figure 1B). In clear plaques, where there is no detected fluorescence, both strains are being lysed by the phage and thus it is likely that the modification made in the second strain is not affecting phage infection. However, in colored plaques, where there is fluorescent signal from the modified strain, only the wildtype strain is being lysed. In this case, it is likely that the alteration in the modified strain has interrupted the phage infection cycle (Figure 1B). This technique enables rapid disregard of the plaques formed by off-target phages and immediate identification of target phage plaques, greatly reducing laborious downstream characterization.

The DisCo method can be further multiplexed by adding more than two strains to the screening lawn. To increase screening throughput, three or more fluorescently-tagged uniquely modified strains can be combined in a screening plate along with an environmental sample, limited by the number of distinguishable detection channels. As before, a clear plaque without fluorescent signal represents a phage that lyses all strains and thus likely does not interact with any of the modifications made to the bacterial strains. However, in this multiplexed design, there is not just one possible fluorescent plaque. Instead, there are multiple possibilities, most importantly single-color plaques in which the modification of the surviving tagged strain impedes phage infection (Figure 1C). This approach allows the detection of multiple orthogonal phage traits in a single one-step screen. Note that if non-orthogonal traits are targeted (i.e. primary & secondary receptors), bi-color plaques can be detected. The extra information gained from co-culturing multiple strains has the potential to speed up targeted phage discovery by orders of magnitude while retaining many of the advantageous features of conventional methodology.

### Phage DisCo can recover characterized phages based on their known receptors

To validate that Phage DisCo works as expected, we first tested the co-culture set up with a pool of previously characterized phages and *E. coli* knockout strains corresponding to each phage’s receptor protein (Supp. Table 1). We imaged each channel using our custom fluorescent plate imager (Supp. Figure 1, Supp. Table 2) and made a composite image. Each phage can be immediately identified by plaque color (Supp. Figure 2). We note that there can be an apparent increase in fluorescence intensity within colored plaques relative to the mixed strain lawn due to the relative increase in single strain density when other strains are unable to grow, and this effect makes single-color plaques particularly easy to detect in composite images. Having established that Phage DisCo can distinguish between well-characterized phages, we proceeded to test our system with environmental samples containing unknown phages.

### Phage DisCo can efficiently isolate novel phages with specific receptor dependencies

To determine how our screen would work with unknown environmental phages, we used the same set of three strains used for the proof-of-concept experiment (Supp. Table 1) and added filtered samples of wastewater from the Boston metropolitan area (MA, USA). As with the characterized phages, after making composite images of all three fluorescent channels we could immediately identify phages putatively dependent on each protein (Figure 2A). To validate the receptor dependency of the wild phages identified in our DisCo screen, we picked a plaque of each color for further downstream processing. We note that without the extra layer of information provided by the fluorescent signal, many of these plaques have similar morphology making them impossible to distinguish as unique. After purification and replication of each phage, we plated each on monoculture lawns of wildtype, knockout, and complemented knockout strains to validate receptor-dependency (Figure 2B, raw data in Supp. Figure 3). As expected, each phage dropped below detection when plated on a receptor knockout strain, and plaquing was fully restored by plasmid complement, validating our DisCo approach for receptor-guided phage discovery.

**Figure 2:**
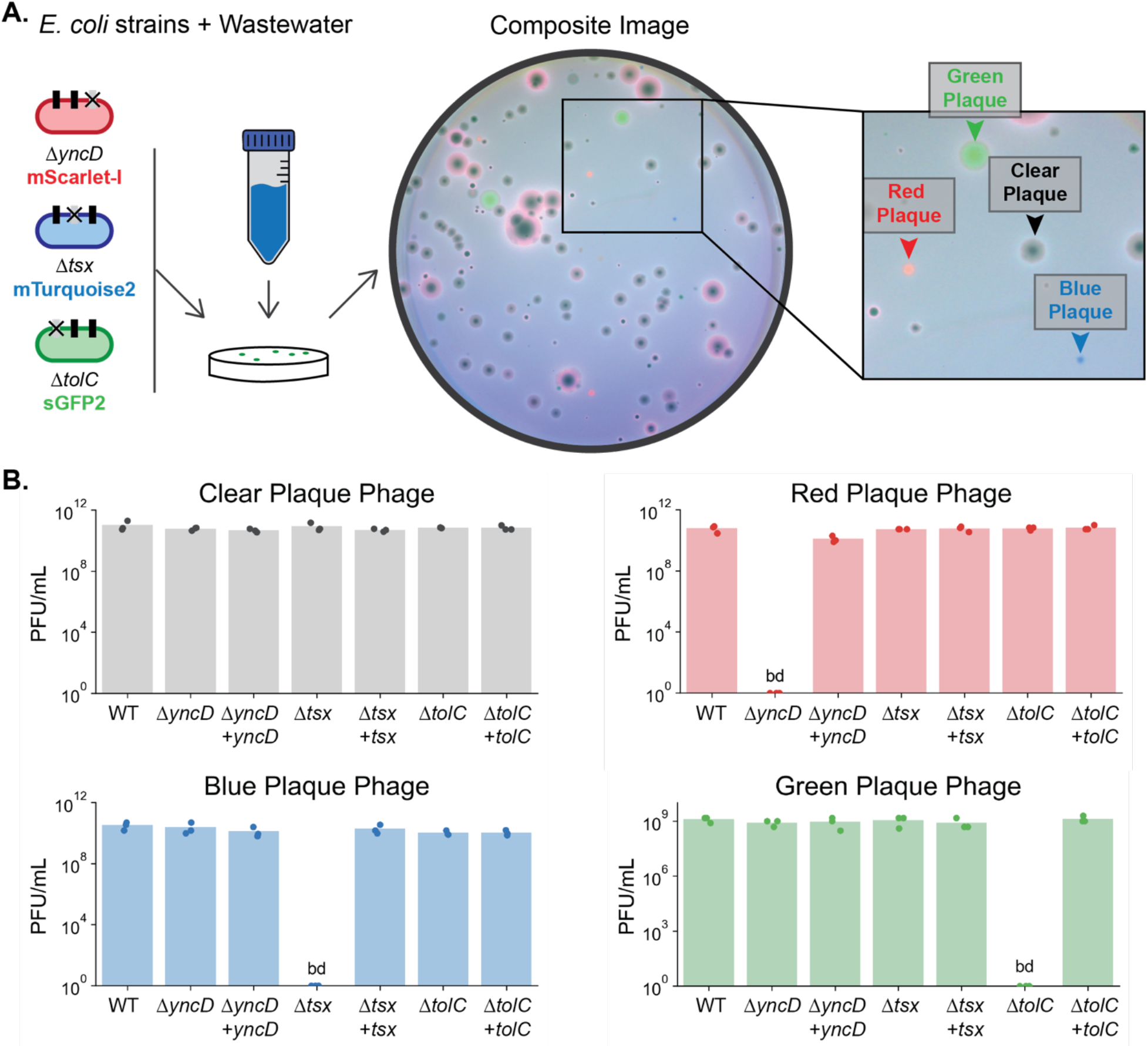
Phage DisCo can discover receptor specific phages. **A**. Screening plate with three *E. coli* knockout strains and a wastewater sample. Arrows point to the plaques selected for analysis. **B**. Each phage was purified, replicated, and three biological replicates were plated for PFU on seven monoculture bacterial lawns. The lawns were wildtype (WT), each of the three knockout strains, and each of the three knockout strains plus a plasmid complement. Counts below detection are noted bd.

To quantify the improved efficiency of using Phage DisCo to identify target-specific phages compared to traditional methods, we recorded the frequency of positive plaques for each receptor target. Of 746 total plaque screened across 5 replicates, 11 were TolC-dependent, 46 were Tsx-dependent, and 11 were YncD-dependent. Given that picking plaques at random to find one dependent on a receptor of interest would follow a negative binomial distribution, it would be necessary to screen on average 67, 15, and 67 plaques before finding a TolC, Tsx, and YncD phage respectively without the extra information provided by Phage DisCo. This result highlights that Phage DisCo not only provides receptor information, reducing laborious downstream characterization, but also that isolation efficiency of rare phages can be improved by over an order of magnitude.

### Discovery of phages interacting with defense systems

Though receptor-specificity dictates the initial outcome of phage infection, it is increasingly recognized that many other bacterial factors can affect the outcome of phage infection [20]. We used Phage DisCo to discover phages that interact with phage-defense systems. We used two isogenic strains per plate where each has a fluorescent tag and one strain is expressing a known phage defense system. If a phage is sensitive to the defense system in question, the defense-negative strain will be fully lysed while the defense-positive strain will continue to grow and emit fluorescent signals, leading to detectable colored plaques. To test this screening method, we conducted a positive control using the phage defense system GmrSD, a modification dependent restriction enzyme which cleaves DNA at the 5hmC modification found commonly in T even phages [21] (Supp. Table 3). Phage T4 is natively insensitive to GmrSD because of the anti-defense internal protein IPI which inhibits the effect of GmrSD [22]. We tested whether phage DisCo could distinguish between GmrSD sensitive and resistant phages (T4 ΔIPI and T4 WT respectively).

As anticipated, T4 ΔIPI and T4 WT plated differentially on the co-cultured lawns (Figure 3A,B) indicating that Phage DisCo plates can identify phages that interact with a specified phage defense system. No fluorescent signal was detected in plaques made by phage T4, but red fluorescent signal was visible where cells expressing GmrSD had resisted infection by phage T4 ΔIPI. In contrast to the phage DisCo screens for receptor-dependent phages, these plaques were associated with a unique fluorescent signal that was intense around the periphery of the plaque but depleted in the center, creating a halo effect. This effect is likely to be caused by the spatiotemporal dynamics of phage concentration across the plaque, and we speculate that the GmrSD defense system may be overwhelmed at high concentrations of phage, as has been previously observed [23]. Having established the utility of the assay to detect GmrSD sensitive and resistant phages, we next used the same assay to screen for wild phages in wastewater samples. Consistently, we were able to rapidly identify plaques with the same fluorescent halo morphology, and we confirmed that the identified phages were sensitive to GmrSD by purifying and plating them on monoculture lawns (Figure 3C/D). Our isolation of wild GmrSD-sensitive phages demonstrates the potential for Phage Disco to identify phages that are interacting with restriction modification-based defense systems, which is an example of a defense system that presumably saves the life of the infected cell. However, recently it has become clear that a large fraction of all known phage defense systems operate via abortive infection, wherein the infected cell dies before progeny phages can be released [24,25]. We reasoned that such mechanisms may be difficult to incorporate into the phage DisCo screening method because successful phage replication in the defense-negative strain would kill the co-cultured defense-positive strain by abortive infection, potentially preventing differential plaquing.

**Figure 3:**
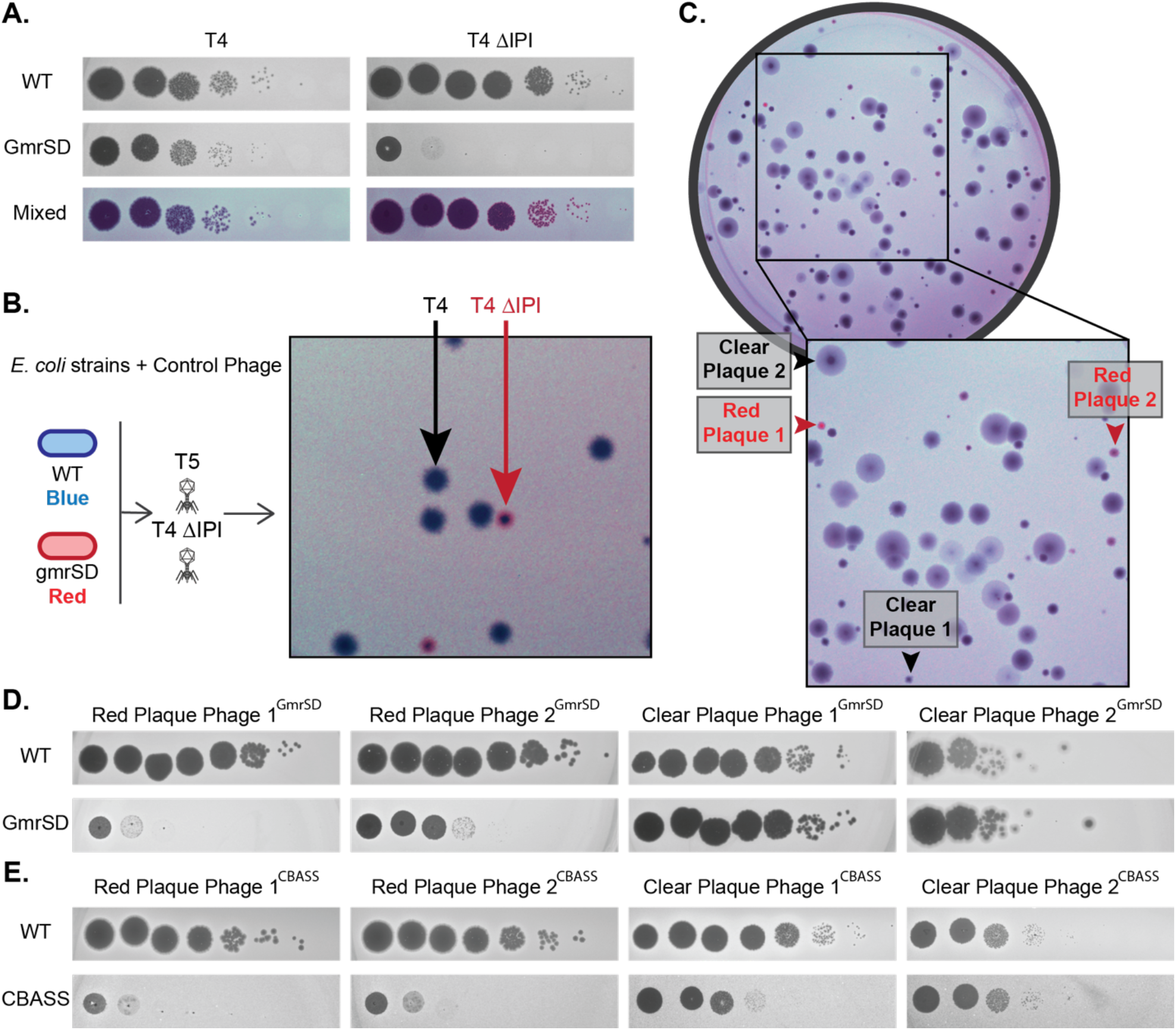
Phage DisCo can discover phages that interact with specific phage defense systems. **A**. Monoculture and mixed lawns with control phages to indicate expected plaque morphology when screening for phage defense system sensitive phages. **B**. Plaque morphology of controls on a simulated Phage DisCo screening plate with known phages. **C**. Wastewater screening plate with the same strains shown in panel A. Two hits are labeled as Red Plaque 1 and 2, and two non-hits are labeled Clear Plaque 1 and 2. **D**. Each of the phages in the plaques highlighted in panel B after purification and replication on monoculture wildtype (WT) and GmrSD containing lawns. **E**. Phages picked from a similar screen for phages that interact with a CBASS system. The screening plate for this assay is shown in Supp. Figure 4.

To test if the Phage DisCo screening approach would work for a known abortive phage defense system, we tested a recently characterized cyclic oligonucleotide-based signaling system (CBASS). Following infection with phages, CBASS systems cause rapid cell death via activation of toxic effector proteins [26]. To test if phage DisCo was able to detect CBASS-sensitive phages, we used a CBASS system capable of defending against a broad range of phages [27]. Despite the potential loss of signal from abortive infection, many putative CBASS-sensitive phages were detected in wastewater samples (Supplemental Figure 4). Two positive and two negative plaques were purified and screened in monoculture to validate the DisCo screen. Both putative CBASS-sensitive phages showed reduced efficiency of plating on strains containing the defense system in monoculture, validating the screen. However, we note that one of the putatively defense-resistant control phages (Clear Plaque Phage 1^CBASS^) also showed lower plating efficiency on the CBASS containing strain as compared to the wildtype strain, implying false negatives may occur with some defense systems (Figure 3E). While we were able to detect phages sensitive to CBASS with our phage DisCo assay, sensitivity may be reduced for abortive infection systems, resulting in decreased ability to detect phages sensitive to this type of defense system.

As a final test of Phage DisCo, we screened for phages that interact with the abortive phage defense system BstA in *Salmonella enterica* [28]. While we were able to validate that Phage DisCo can determine the difference between phages sensitive to BstA including temperate phage BTP1 (Supp. Figure 5), no BstA-sensitive environmental phages were detected in our screens. This suggests that BstA-sensitive phages may be rare in the environmental samples screened, though we cannot rule out the possibility of false negatives, as observed for the CBASS system.

## Conclusion and Discussion

In this paper, we have expanded upon the widely used traditional plaque assay for phage screening and isolation by co-culturing multiple fluorescently-tagged bacterial host strains in a single lawn to create an efficient, targeted assay. We validated this assay by testing previously characterized phages and showed that each phage was distinguishable by its known receptor. Our experiments also showed that we could identify phages from environmental samples with desired characteristics in a single screening step, which drastically cuts down on the steps and time required by current techniques. We also validated that the method can distinguish between control and environmental phages that interact with a given defense system by slightly altering the Phage DisCo configuration. Based on work presented here and previous published work [17], we conclude that Phage DisCo can substantially reduce time and labor needed for targeted discovery of lytic or lysogenic phages with DNA or RNA genomes.

We can see possible applications for Phage DisCo across multiple fields. Targeted screening methods can help survey ecological relationships across environments and within the same environment over time. Understanding phages within an environment could help us understand some of the selective pressures on bacteria, for example what defense systems may be beneficial or redundant in contemporaneous bacterial populations. Screening for differential plaquing across strains can also give insights into interactions between host genes such as receptors and phage infections. This can be utilized to quickly compile diverse phage cocktails for phage therapy as using phages with diverse receptors within a phage cocktail is thought to avoid easy pathways to resistance [29]. Similarly, there have been suggestions to ensure that all phages in a cocktail are not all sensitive to the same defense system [30]. Phage DisCo can quickly survey an environment for both axes of diversity and could even be nested to find phages with very specific properties, and may even be applicable to the discovery and culture of eukaryotic viruses

Here we have shown receptor knockouts and defense system knock-ins in lab strains of *E. coli*, but modifying environmental strains may help discover more environmentally relevant phages. For example, instead of heterologously expressing a defense system in *E. coli* wildtype bacterial isolates containing defense systems together with isogenic knockouts could allow more physiologically-relevant screens, potentially increasing positive hit rate. Furthermore, the use of Phage DisCo to isolate defense system-sensitive phages has the potential to provide insights into the molecular mechanisms of these systems themselves, as the evolution of resistant “escape” phages can pin-point phages proteins that trigger their activity [31].

Although we anticipate Phage DisCo will be generally applicable to targeted phage discovery efforts across many bacterial study systems, there are limitations. To set up a Phage DisCo screen, genetic tractability is required to introduce fluorescent tags and genetic perturbations between the isogenic bacterial strains. Additionally, in the case of multiplexed screens using multiple knockout strains on the same plate (Figure 1C), we assume that the proteins knocked out in the three strains are independent of each other within the bacterial host and during phage infection. If this is not the case, and the proteins are in some way dependent on each other, a phage that depends on any one of them will fail to make a plaque at all and false negatives can occur. Replacing one knockout strain with a wildtype strain can reduce such false negatives at a slight cost to throughput. Similarly, care must be taken when interpreting data from Phage DisCo assays biologically: phages frequently have multiple receptors, which often include genetically complex structures such as lipopolysaccharides. Phage DisCo only shows the impact of a single genetic perturbation on phage infection, and perturbation of genes associated with epistatic effects (e.g. genes in the LPS biosynthetic pathway) may be difficult to interpret in terms of phage biology. In this case, follow up, for example using the INSeq method [8], would be needed to verify the full dependencies of any given phage.

Phage DisCo has the potential to rapidly rule out common phages in any given sample and highlight rare phages with desired characteristics, improving the efficiency of culture-based virus discovery. We believe this method will vastly improve phage prospecting and help lead to discoveries in phage research.

## Materials and Methods

### Bacterial strains, plasmids, and phages

*E. coli* knockout strains (Δ*yncD*, Δ*tsx*, and Δ*tolC*) were taken from the Keio collection [32] and the complement plasmids were taken from the ASKA collection [33]. Correct identity of the Keio Strains were confirmed using PCR amplification of the knockout region followed by sequencing, and AKSA strains were confirmed by extraction with the NEB Monarch Plasmid Miniprep Kit followed by sequencing. The pEB2 plasmids were adapted from Addgene #104007 to change kanamycin resistance (KanR) to chloramphenicol resistance (CmR) which is compatible with the Keio strains using overlap extension PCR cloning to construct the final plasmid [34,35]. The mScarlet-I fluorescent protein was switched for mTurqoise2 and sGFP2 using the same method. The pEB2 plasmids were pooled together and then transformed into the Keio strains by electroporation. Briefly, strains were grown in LB broth without salt (10g/L tryptone, 5g/L yeast extract) until optical density (OD) ∼0.5, washed in cold, sterile water and electroporated at a resistance of 200 Ohms, capacitance of 25μFD, and 1.6 Volts. Transformants were selected for on 2% LB agar plates + 20 μg/mL chloramphenicol. GmrSD was amplified out of MRSN781659 cloned into the low copy pSC plasmid backbone using NEBuilder® HiFi DNA Assembly Master Mix. The pBR322 plasmids were created by modifying the NEB vector (catalog #N3033S). (Summary of the plasmids used and modifications made is in Supp. Table 4.) All strains were grown in LB Lennox broth (10g/L tryptone, 5g/L yeast extract, 5g/L NaCl). ASKA plasmids were selected for with 20 μg/mL chloramphenicol. No selection was required for the pEB2 or pBR322 plasmids after cloning.

### Wastewater collection and processing

Primary effluent wastewater samples were collected from two separate sites near Boston, MA, USA. Samples were centrifuged at 4000 X g for 30 minutes, and the supernatant was filtered through a 0.22μm vacuum filter. Filtered wastewater was stored at 4°C until use. Classical double-layer plaque assays were used to titer the total detectable phages in each sample by adding 10, 100, and 300μL wastewater to 100uL of saturated wildtype *E. coli* BW25113 culture. The cells and wastewater were combined with 3mL of LB Lennox top agar (LB Lennox media + 0.5% Agar) at 55°C and poured onto a solidified LB + 2% agar plate. Plates were incubated at 37°C overnight to enumerate phage titers. Optimal plates have ∼150 plaques per 10 cm petri dish, this is approximately the maximum number that can be screened per plate without frequent overlap between plaques.

### Phage DisCo plaque assay

For the receptor screen, colonies of each fluorescently tagged Keio strain were added to 2mL of LB Lennox broth without selection and grown until the culture was saturated. 100μL of each culture was added to 100-300μL of filtered wastewater sample. Volume of wastewater was optimized based on total detectable phage concentration by the titer method described above and varied slightly depending on wastewater sample date and location. To make each screening plate, 3mL of LB Lennox top agar at 55°C was added to the cells and wastewater mixture and poured onto a 10 cm petri dish with solidified LB + 2% agar incubated at 37°C overnight.

For the GmrSD screening plates, a similar protocol was followed with the addition of arabinose at a final concentration of 0.1% to induce production of the GmrSD protein. Likewise, arabinose was added to induce the production of the CBASS construct and anhydrotetracycline (AHT) was added at a final concentration of 500 ng/mL to induce the production of the BstA construct [28].

### Fluorescent imaging

After overnight growth, Phage DisCo plates were imaged in our custom fluorescent plate imager (diagram and parts list in Supp. Figure 1 and Supp. Table 2 respectively). In short, the plates are illuminated by colored LEDs with excitation filters (EX), the emitted light passes through emission filters (EM), and images are taken on a Canon EOS camera. The red channel pairs 567 nm LED with 562 nm EX and 641/75 nm EM filters, the green channel pairs 490–515 nm LED with 494 nm EX and 540/50 nm EM filters, and the blue channel pairs 448 nm LED with 438 nm EX and 483/31 nm EM filters (summary of which channels were used to detect which fluorescent signals in Supp. Table 4). Exposure times varied between 0.5s and 8s depending on the number of strains on the plate and intensity of fluorescent signal. Exposure times were selected such that there was little to no fluorescent signal within the plaques in each channel while maintaining bright signal on the rest of the plate. Camera settings were kept at an aperture of 5.6 and ISO of 200.

Once the individual channels were imaged, composite images were created in Adobe Photoshop. Some channels were linearly adjusted for contrast to increase signal to noise. No conclusions were made using fluorescence intensity. Hits are based solely on presence or absence of fluorescent signal.

### Purifying phages from Phage DisCo screening plates

Once hits were identified in composite images, phages were picked from plaques into 200μL of SM buffer (200 mM NaCl_2_, 10 mM MgSO_4_, 50 mM Tris-HCl, pH 7.5). The resuspended plaque was then filtered using 0.22μm spin-x centrifuge tube filters. The filtrate was serially diluted in more SM buffer and then plated on monoculture lawns to confirm the predicted characteristics. These lawns were made by mixing 100μL of cells, and 3mL of LB Lennox Top Agar at 55°C, with arabinose if required, and pouring the mixture onto a solidified 2% LB agar petri dish. 2μL of each dilution was spotted on top of the solidified cell and agar mixture. These spots were allowed to dry and then the plates were incubated at 37°C overnight. Once the phenotype was confirmed, phages were replicated by adding 2μL of phage to 2mL saturated bacterial culture diluted 1:1000 in LB broth. These new cultures were grown overnight at 37°C with aeration then centrifuged at 16,900 X g for 5 minutes and the supernatant passed through a 0.22μm filter to remove bacterial cells. These high titer stocks were used to make another set of dilution plates following the same protocol for figures and stored at 4°C.

### Complement assays

Complement assays were done using the Keio background strain (*E. coli* BW25113), the Keio knockout strains, and Keio knockout strains plus the corresponding ASKA expression plasmids. To make the complemented knockout strains, the corresponding ASKA strains were grown in LB Lennox broth with 20μg/mL chloramphenicol. Plasmids were extracted from the ASKA strains using the NEB Monarch Plasmid Miniprep Kit and transformed into Keio strains by electroporation as above. To test the plaquing ability of each phage, three sets of serial dilutions were made to provide biological replicates. The dilutions were arrayed in a 96 well plate, and lawns were made using each strain on rectangular Omnitrays (Nunc). Each lawn had 100μL of cells and 5mL of LB Lennox Top Agar at 55°C. Plates for the Δ*tsx*+*tsx* and Δ*yncD*+*yncD* strains were made using 20μg/mL chloramphenicol in the 2% LB agar. 2μL of each of the dilutions was spotted onto each lawn using a Gilson Platemaster (96 well pipette). Spots were allowed to dry and then plates were placed in plastic bags to prevent drying of the lawns during the overnight incubation at 37°C. Plates were then scanned using an Epson Perfection V850 Pro Scanner. The dilutions in the raw data shown in Supp. Figure 3 represent the -3 through -10 dilutions.

### Example Protocol

#### Materials required

- Bacterial strains
  - Fluorescently tagged either chromosomally or on a plasmid
    - Note: Fluorescent tags must be constitutively expressed, and we found the extra signal provided by multicopy plasmids as compared to a single copy on the chromosome to be helpful
  - At least one strain with a genetic perturbation expected to impact phage infection
- Media and agar
  - Media for the chosen bacterial host
  - Petri dishes containing media + solidified 2% agar
  - Molten media + 0.5% agar (top agar)

#### Equipment required

- Fluorescent imager
  - Capable of imaging full dishes at macroscopic level
  - Filters compatible with chosen fluorescent tags

#### Protocol

1. Inoculate one colony of each strain into individual aliquots of media and allow to grow until saturated. Time, temperature, aeration, etc. will depend on the bacterial host.
2. Once grown, mix equal volumes of each strain for each DisCo plate.
  - Note: Volume needed to make a smooth lawn of bacteria will depend on the bacterial host. For lab *E. coli* strains used here, 100uL of each strain was added to each plate.
3. Add filtered environmental sample to cell mixture.
  - Note: Volume of environmental sample will depend on concentration of phages. See Wastewater Collection and processing section above for titer methods.
4. Add enough of the molten top agar to cover the screening dishes.
  - Note: For 10 cm petri dishes, 3mL of top agar is recommended. For 128 × 86mm rectangular dishes (SBS sized plate), 5mL of top agar is recommended.
5. Pour the cell, sample, and top agar mixture onto the solidified 2% agar dish being careful not to add any bubbles.
6. Allow to set at room temp for ∼2 minutes.
7. Incubate overnight at the correct temperature for optimal growth of the chosen bacterial strain.
  - Note: Incubating inside an airtight plastic bag can help reduce drying for longer incubations and make images more even.
8. The next day, remove from the incubator and image each fluorescent channel individually.
9. Combine the channels into one image to detect differential plaquing.

When using lab *E. coli* strains, equal volumes of each strain were added to each screening plate without normalization for optical density or CFU. Even when strains grew at different rates, differential screening was detectable as a consequence of strains growing denser (and brighter) within partial plaques. This is particularly true if including a wildtype strain on a two strain DisCo plate or if including three orthogonally perturbed strains on a DisCo plate. If using the Phage DisCo method under different conditions or with another bacterial host, it may become more important to control for these factors and others such as log phase growth. If lawns look patchy or grainy or if signals are hard to detect, it is recommended to grow cells to log phase and then concentrate by centrifuging the liquid culture at 16,900 X g for 5 minutes, removing the supernatant, and resuspending in 10x less volume. After concentrating, continue with the method as written above.

## Acknowledgements

We thank Lucy McCully, Fernando Rossine, and Kepler Mears and the rest of the Baym lab for helpful discussions and materials. We thank Daniel Eaton and Johan Paulsson for making both the Keio and ASKA collections available to us, Alita Burmeister and Paul Turner for sharing a sample of phage U136B, Akos Nyerges and George Church for providing phage samples, and Peter Wiegele and Rebekah Silva at NEB for the modified T4ΔIPI phage. We thank Sam Hobbs, Desmond Richmond-Buccola and Philip Kranzusch for useful discussions and providing resources. We thank the staff at the Multidrug-Resistant Organism Repository and Surveillance Network (MRSN) for providing strain MRSN 781659. The custom fluorescent imager was built using tools and with assistance from the Research Instrumentation Core at Harvard Medical School. This work was supported by the NIGMS of the National Institutes of Health (R35GM133700), the David and Lucile Packard Foundation, the Pew Charitable Trusts, Alfred P. Sloan Foundation and NSF grant IOS-2331228. EAR acknowledges support from the Systems, Synthetic, and Quantitative Biology PhD program training award (T32GM135014) and from NIH NIAID F31 (F31AI178993).

