## Supplemental Tables and Figures for "Phage DisCo: targeted discovery of bacteriophages by co-culture"

***Supplemental Table 1:*** Characterized phages and *E. coli* strains used to validate Phage DisCo

| **Characterized phages** | |
|---|---|
| **Phage** | **Receptor** |
| T4 | OmpC |
| U136B [10] | TolC |
| Bas37 [36] | Tsx |
| Bas 10 [36] | YncD |
| ***E. coli* BW25113 strains** [32] | |
| **Modification** | **Plasmid** |
| $\Delta$*tolC* | pEB2-chlor-sGFP2 |
| $\Delta$*tsx* | pEB2-chlor-mTurquoise2 |
| $\Delta$*yncD* | pEB2-chlor-mScarlet-I |

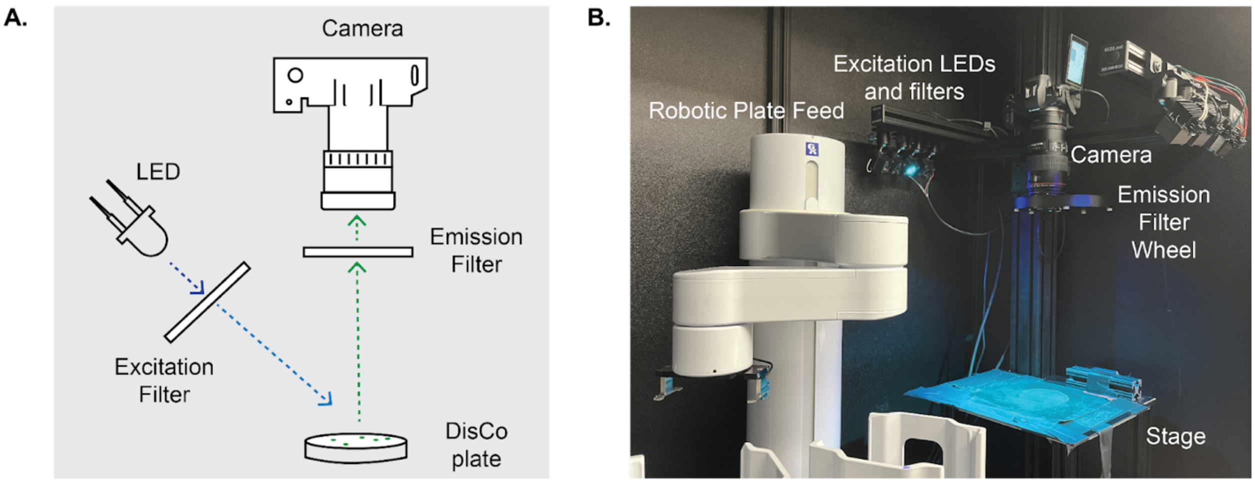


**Supplemental Figure 1: *Fluorescent imaging set up.* A.** Conceptual image of the bare bones needed for any macroscopic fluorescent imager. **B.** Image of custom set up with an array of excitation LEDs and filters, an emission filter wheel, and a robotic plate feed for high throughput imaging. Parts list for our imager in Supplemental Table 2.

***Supplemental Table 2:*** Parts list for custom fluorescence imager

| **Fluorescence** | | | |
|---|---|---|---|
| **Channel** | **Excitation LED** | **Excitation Filter** | **Emission Filter**<br>50mm diameter<br>Edmund optics<br>Format - X/Y<br>X - center wavelength (nm)<br>Y - width of band (nm) |
| Red | Lime (567nm) | 562nm | 641/75 |
| Blue | Royal blue (448nm) | 438nm | 483/31 |
| Green | Cyan (505nm) | 494nm | 540/50 |
| **Camera** | | | |
| Canon EOS R | | | |
| **Lens** | | | |
| Canon EF 100mm f2.8 Macro IS USM Lens | | | |

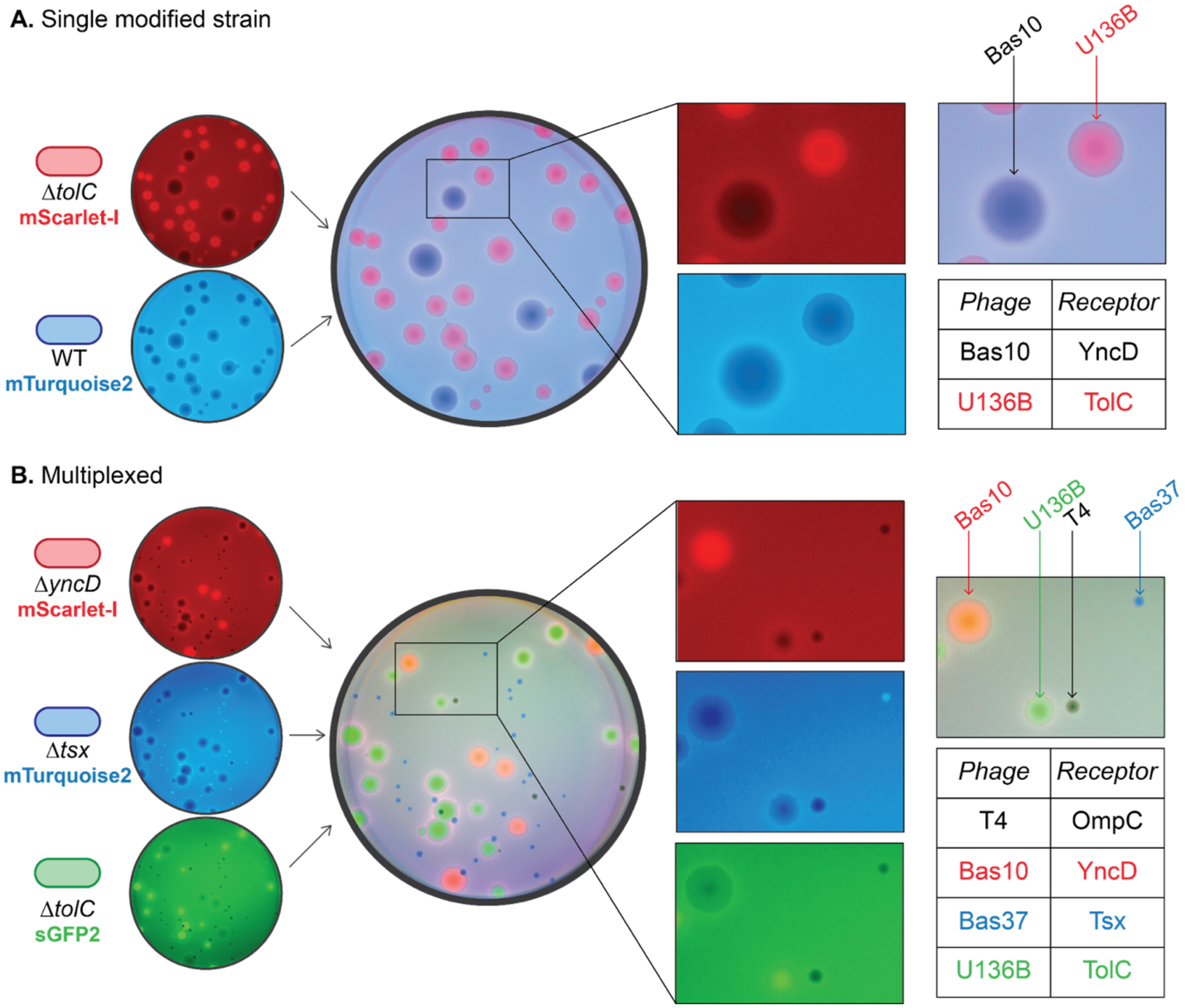


**Supplemental Figure 2:** ***Positive controls show expected plaque morphology on Phage DisCo plates*. A.** Two strain plate with a wildtype (WT) E. coli strain tagged blue and a tolC knockout strain (Δ*tolC*) strain tagged red. As expected, Bas 10 plaques are clear (no fluorescent signal) while U136B plaques are red (red fluorescent signal) where the Δ*tolC* strain continues to grow. **B.** Multiplexed plate with three strains showing that, in this case, no WT strain is necessary and we can detect all three colored plaques. Phage T4, which does not require any of the individually knocked-out proteins for infection, lyses all three strains and leaves a dark plaque with no fluorescent signal. Phage U136B, which has been characterized to require the efflux protein TolC for infection [10], lyses all but the tolC knockout strain. Within the U136B plaques, the ΔtolC strain (tagged with GFP) alone is able to grow and therefore these plaques appear as green in the composite image. Similarly, phage Bas 37, which requires the Tsx protein, has a blue fluorescent signal corresponding to the Δtsx strain, and phage Bas 10 which requires YncD and has a red fluorescent signal corresponding to ΔyncD [20].

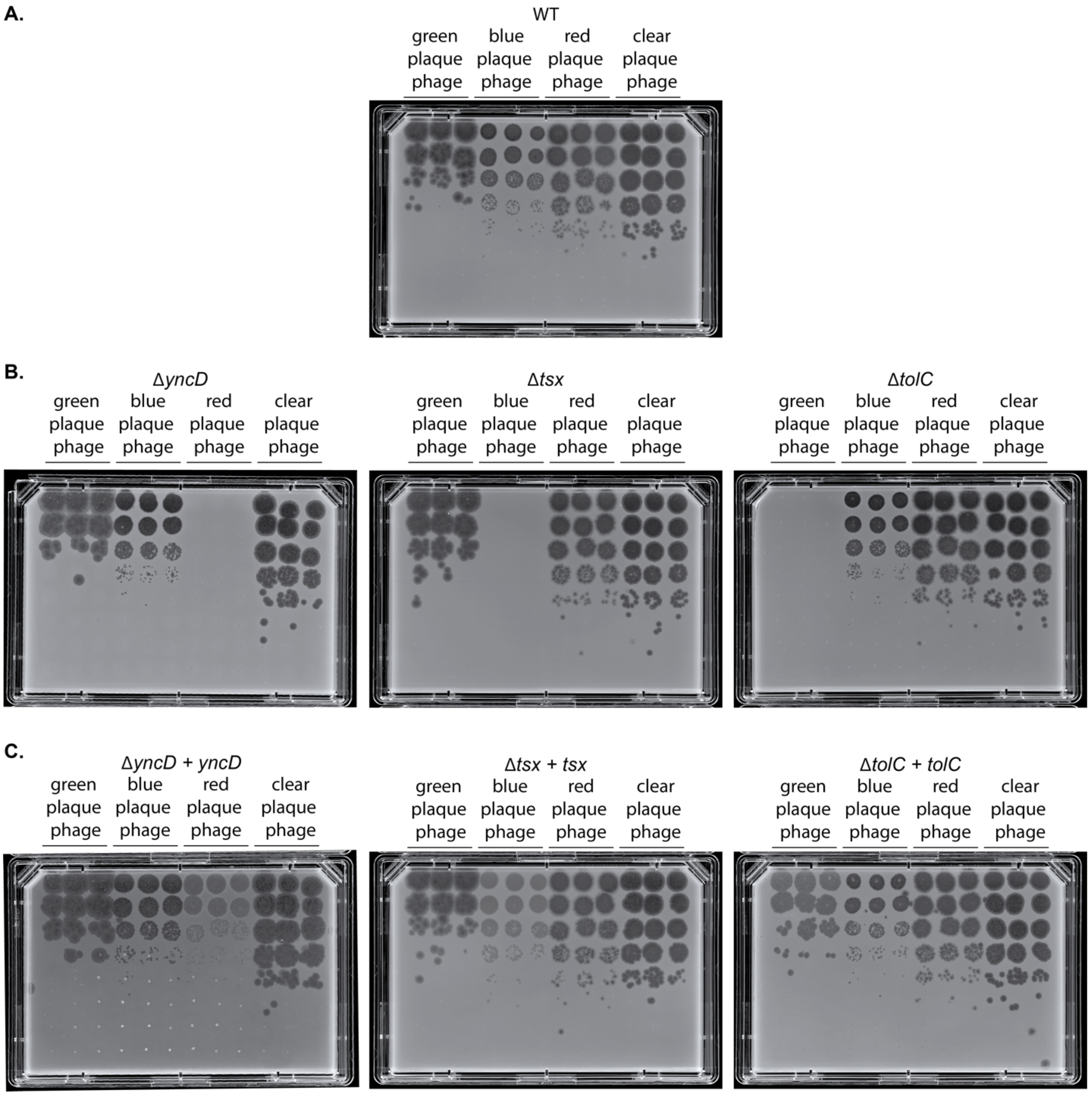


**Supplemental Figure 3:** ***Plaque assays of each phage picked from Figure 2A plated on all seven potential hosts in monoculture which were used to create graphs in Figure 2B.*** **A.** All phages plated on the wildtype (WT) lawn. **B.** All phages plated on each of the knockout strains (Δ*yncD*, Δ*tsx*, and Δ*tolC*) in monoculture. **C.** All phages plated on each of the complement strains (Δ*yncD* + *yncD*, Δ*tsx* + *tsx*, and Δ*tolC* + *tolC*) in monoculture. For all plates, three separate serial dilutions of each phage were made to provide biological replicates. 2uL of each dilution was spotted on each plate using a Gilson Platemaster (96 well pipette). The dilutions shown represent the -3 through the -10 dilutions.

***Supplemental Table 3:*** Phages and *E. coli* strains used to validate phage defense Phage DisCo

| | |
|---|---|
| **GmrSD Experiments** | |
| **Characterized phages** | |
| **Phage** | **GmrSD susceptibility** |
| T4 | No |
| T4 ΔIPI [22] | Yes |
| ***E. coli* MG1655 strains** | |
| **Defense system** | **Plasmid** |
| - | pBR322-mWatermelon |
| pSC-gmrSD | pBR322-mScarlet-I |
| **CBASS Experiments** | |
| ***E. coli* MG1655 strains** | |
| - | pEB2-chlor-mTurquoise2 |
| pBADS-CBASS_EcCdnD_4TM (Addgene #224395) | pEB2-chlor-mScarlet-I |
| **BstA Experiments** | |
| ***Salmonella enterica* LT2 strains** [28] | |
| BstA-STOP | pEB2-chlor-mTurquoise2 |
| BstA | pEB2-chlor-mScarlet-I |
| **Characterized Phages** | |
| Phage | BstA susceptibility |
| BTP1 | No |
| BTP1Δ*bstA* [28] | Yes |

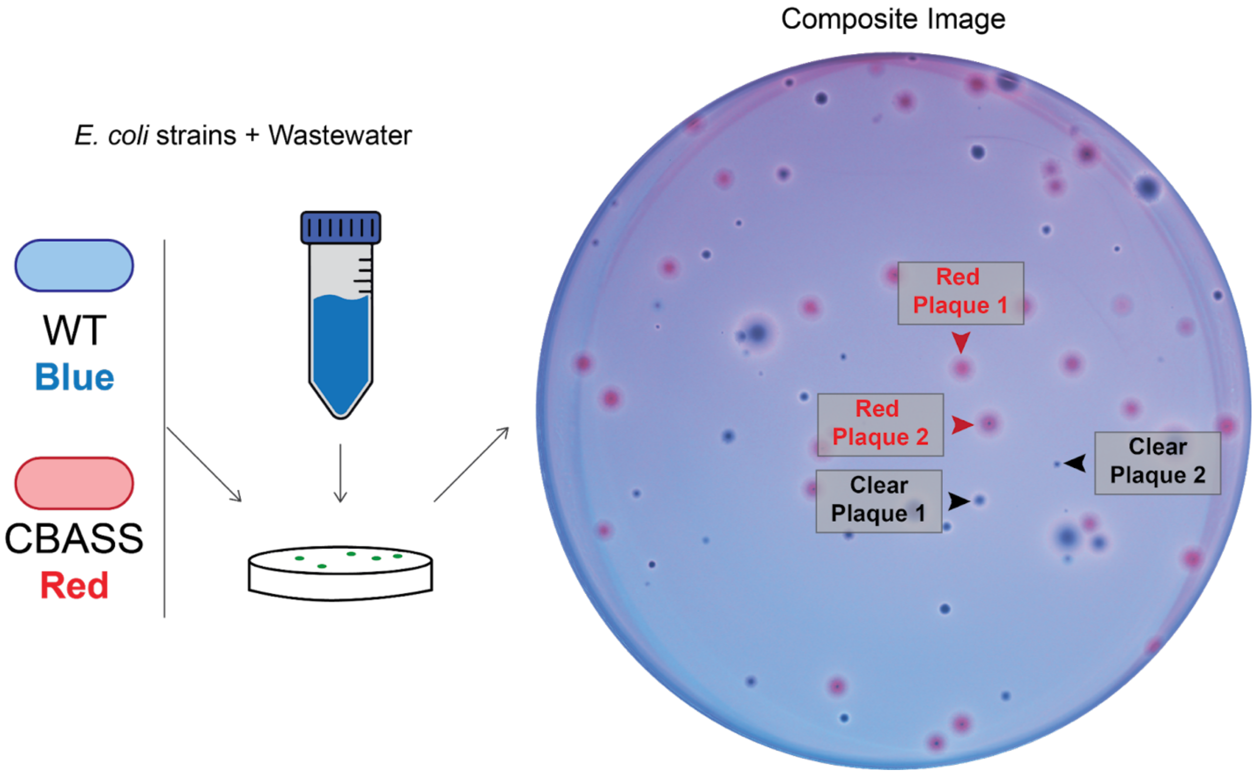


**Supplemental Figure 4:** ***CBASS screening plate shows many hits in wastewater sample.*** Here red plaques are predicted to be interacting with the CBASS defense system and clear plaques are not. The highlighted plaques were picked, replicated, and plated on monoculture lawns to make the images shown in Figure 3E.

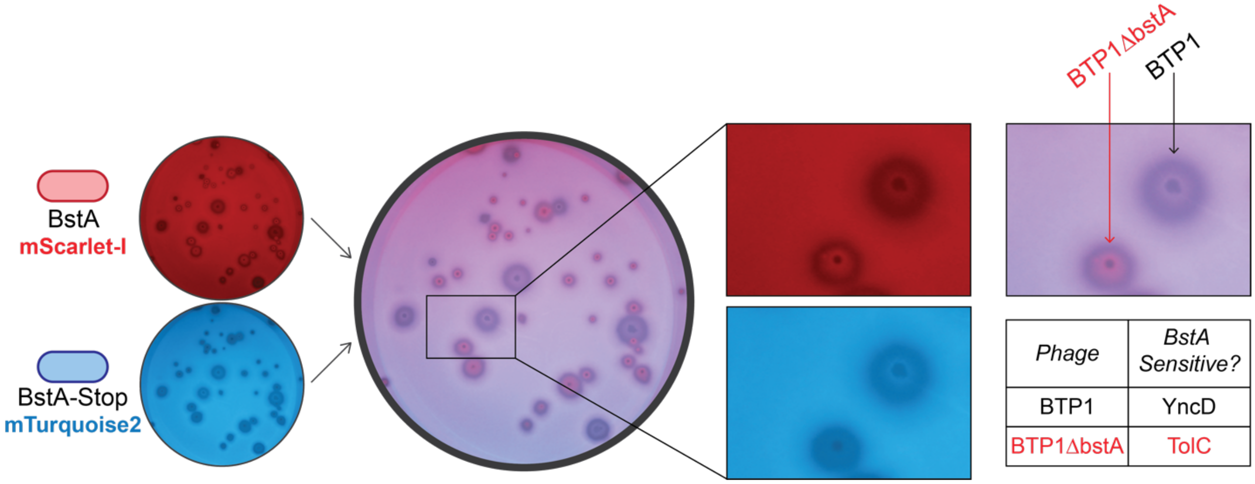


**Supplemental Figure 5:** ***BstA control plate shows that lysogenic phages are also detectable using Phage DisCo.*** Shown here is a control plate with two *Salmonella enterica* LT2 strains. BstA, which is encoded by the temperate phage BTP1, acts as a defense system against phages invading the bacterial host in which it has integrated. However, when BTP1excises from the chromosome to lyse its host cell, it has an anti-defense region that allows for its life cycle to continue unimpeded. For this reason, BTP1is expected to be able to lyse a cell with BstA on the chromosome while BTP1Δ*bstA* is not [28]. The blue strain contains a BstA sequence with an early stop codon while the red contains a functioning BstA defense system. As is anticipated, BTP1makes a clear plaque while BTP1Δ*bstA* has red fluorescent signal within the plaque.

***Supplemental Table 4:*** Summary of plasmids used in Phage DisCo experiments and fluorescent detection conditions for each one

| Plasmid | Description | Selectable Marker | Filters Used for Detection (EX-EM) |
|---|---|---|---|
| pEB2-chlor-sGFP2 | Derivative of Addgene #104007 where Switch KanR for CmR and mScarlet-I for sGFP2 [37] | CmR | 494nm – 540/50nm |
| pEB2-chlor-mTurquoise2 | Derivative of Addgene #104007 where Switch KanR for CmR and mScarlet-I for mTurquoise2 [37] | CmR | 438nm – 483/31nm |
| pEB2-chlor-mScarlet-I | Derivative of Addgene #104007 where Switch KanR for CmR | CmR | 562nm – 641/75nm |
| pBR322-mWatermelon | ORI and rop from NEB #N3033S with KanR and mWatermelon [38] | KanR | 494nm – 540/50nm |
| pBR322-mScarlet-I | ORI and rop from NEB #N3033S with KanR and mScarlet-I [37] | KanR | 562nm – 641/75nm |
